# Repeated environmental changes mitigate inter-individual inequality of food gain in a group foraging experiment with large-billed crows

**DOI:** 10.1101/2025.02.12.637811

**Authors:** Yasuyuki Niki, Mayu Takano, Ei-Ichi Izawa

## Abstract

Animal group foraging produces inequality of food gain among individuals due to their differences, particularly dominance rank. External environmental factors, such as the spatial distribution of food, may mitigate the influence of social features on individual food acquisition. However, the role of environmental fluctuation in social foraging inequality remains largely unexplored Here, we investigated how repeated environmental changes affect inequality in food acquisition among captive large-billed crows (*Corvus macrorhynchos*). By repeatedly altering the relationship between feeding sites and food quantity, we examined changes in the inequality of food gains using Gini coefficient. In the short-term, food allocation became unequal immediately after each change but subsequently recovered. Over successive fluctuations, inequality in food acquisition declined in the long-term, while group foraging efficiency increased. We observed a significant influence of sex on food acquisition, while the expected effect of dominance rank was not detected. Our findings provide the first experimental evidence that changing environments can mitigate inequalities in food gains within foraging groups, highlighting the importance of incorporating naturalistic environmental changes into experimental designs for future group-foraging studies.

## 1. Introduction

Inequality is a defining feature of animal societies, including human society [1–3]. While animal groupings benefit individuals by efficiently finding potential predators [4,5] and food resources [6], they cost through intensified competition for scarce resources. Competition for habitat, mating opportunities, and food resources leads to unequal resource acquisition among group members due to difference in social attributes and phenotypes, including kinship, age, sex, and dominance rank [7].

In group foraging animals, unequal food acquisition occurs among individuals because group members compete for resources that are both spatially and temporally uncertain and finite [8,9]. Dominance rank, based on an individual’s resource-holding potential (RHP), represents a key factor contributing to inter-individual inequality in food acquisition [10,11]. Individual variations in RHP determine access priority to resources, leading to the establishment of dyadic dominant-subordinate relationships and dominance rankings within a group.

Both theoretical and empirical studies have explored how dominance influences individual food gains in group-foraging contexts. Game-theoretical models propose a stable evolutionary strategy whereby subordinates adopt a “producers” role, actively exploring and discovering food sources, while dominants employ “scroungers” tactics to exploit the food produced by subordinates [12,13]. This producer-scrounger dynamic has received empirical support across diverse taxa. For instance, field experiment with group-foraging capuchin monkeys (*Cebus apella*) demonstrated that subordinates initially benefit from their “producer” role when discovering new foods. However, this advantage proves transitory as dominants subsequently monopolise these resources upon arrival [14]. These dominance-based foraging tactics consistently result in unequal food gains, with dominants securing greater overall food intake than subordinates. Similar patterns have been documented across group-foraging bird species including geese (*Branta leucopsis* [15]) and multiple corvids species (ravens, *Corvus corax* [16]; northwestern crows, *Corvus caurinus* [17]; Mexican jays, *Aphelocoma ultramarina* [18]).

Individual differences in food acquisition result from both intrinsic factors (such as dominance rank) and extrinsic environmental factors, particularly the spatial distribution of food resources [19,20]. Experimental evidence demonstrates that environmental context can modulate dominance effects on foraging success. For example, experiments with ruddy turnstones (*Arenaria interpres*) revealed that dominants interfered with subordinates and secured more food when resources were spatially concentrated, but these advantages disappeared when identical food quantities were dispersed throughout the experimental area [21]. This finding highlights the importance of considering environmental factors alongside social hierarchy when examining foraging inequality. Despite these insights, the role of environmental variability in mediating social foraging dynamics has received little attention in the literature, representing a major knowledge gap.

Moreover, the environmental changes, such as spatial distribution of food, over time remains poorly understood in group foraging contexts. Natural foraging environments are inherently variable, with food resource availability fluctuating due to seasonal changes in patch quality, variation in accessible food types [22,23]. Even in the absence of seasonal variation, repeated foraging at sites reduces local food availability, compelling foragers to relocate to alternative patches. Previously depleted sites may subsequently recover due to reduced consumption pressure, creating fluctuating resource opportunities that foragers must continuously decide whether to stay or move on to find food [24]. Such fluctuating conditions may fundamentally alter the costs and benefits associated with dominance-based foraging strategies. Subordinate individuals, typically displaced from prime feeding locations by dominants, may benefit from environmental changes that create new resource opportunities. This potential advantage may stem from subordinates documented greater behavioural flexibility and enhanced learning abilities compared to dominants [25, 26]. Furthermore, temporally variable environments may promote more complex individual strategies and inter-individual interactions, including recursive movement patterns [27] and adaptive changes in foraging motivation driven by environmental uncertainty [28]. Despite these considerations, the role of environmental fluctuation in social foraging inequality remains largely unexplored.

Corvids, particularly crows and ravens are suitable for investigating environmental effects on foraging inequality due to their well-documented group-foraging behaviour [29] and clear expression of producer or scrounger tactics based on their dominance rank, leading to unequal food acquisition [16,17,30]. Large-billed crows (*Corvus macrorhynchos*) are particularly suitable for experimental investigation as they form stable, linear dominance hierarchies in captive groups [31,32], with males typically dominant over females.

In this study, we investigated how repeated environmental changes in the spatial distribution of foods affect inequality in food acquisition and foraging performance within a captive group of large-billed crows. We placed three feeding sites and altered the relationship between each site and their food content, simulating situations in which rich feeding sites become depleted through consumption while previously unused sites recover over time. To focus on food acquisition and foraging performance of group, the total amount of food remained constant throughout the experiment. In this way, we examined the effects of repeatedly changing on a foraging caused by cycles of depletion and recovery.

We evaluated the effect of repeated environmental fluctuations at both group and individual levels. At the group level, we quantified inequality in food aquation within foraging group using Gini coefficient, a well-established measure of resource distribution equality among group members. In addition, we examined the number of individuals successful foragers and time efficiency to search for and consume food. At individual level, we examined how dominance rank and sex influenced food acquisition during environmental changes. As previously reported, dominant individuals may monopolise food in stable situations [14,21]; however, immediately following an environmental change, subordinate individuals may discover and consume food before others. Consequently, inequality in food acquisition at the group level may be reduced. Additionally, sex may alter food preferences and the use of social information [5,33]. Such variations may potentially influence food acquisition in the present group-foraging experiment.

## 2. Materials and Methods

### 2.1 Animals

We used 10 juvenile large-billed crows (5 males and 5 females) aged 2 years, weighing between 640 and 820 g. Sex was identified by analysing DNA from blood samples. The birds were captured as free-floater yearlings in the Yokohama and Kamagaya areas from October to December 2022, with authorisation from Japan’s Ministry of the Environment (license no. 041454056). No individuals were paired during the experimental period. The captured birds were group-housed in an outdoor aviary measuring 5 × 10 × 3 m. The aviary had sufficient wooden perches for all the birds to perch simultaneously, and tubs of water were supplied for drinking and bathing. Slight food deprivation, specifically no food 1 h before the start of daily sessions, was used to encourage the crows to forage in the experimental setting. Except during deprivation periods and experimental trials, food (dried dog food, eggs, and vitamin and mineral supplements) was always available in each compartment.

### 2.2 Ethical note

All experimental protocols and housing conditions followed the Japanese National Regulations for Animal Welfare and were authorised by the Animal Care and Use Committee of the Keio University (No. A2022-348).

### 2.3 Dominance rank assessment

#### 2.3.1 Observations of agonistic interactions

Before starting the group foraging experiment, the dominance ranks of individuals in the group were established using social interaction data collected through daily observations from April to June 2023. Daily observations based on video recording were performed using two omnidirectional video cameras (PIXPRO sp360 4 K; Eastman Kodak Co., Rochester, NY, USA), which were placed at the centre of the aviary ceiling to capture the entire space, including perches and floors, and to allow collection of all social interactions. Video recordings were conducted for 30 min, twice daily, from 8:00–9:00 and 14:00–15:00. A researcher (M. T.) used the behaviour coding software BORIS version 7 [34] to code agonistic interactions offline from video-recorded data. Dominance rank was determined using a total of 8.5 h of video data. We defined an agonistic interaction as a submission display (submissive begging) where the recipient avoids or retreats from the antagonist’s aggressive behaviour. Aggressive behaviours included pecking, kicking, and displacement techniques, while submissive behaviours include avoiding and retreating [31,32]. In each agonistic interaction, the bird that demonstrated submission, avoidance, or retreat was deemed the loser, whereas the other bird was the winner. We noted the winners and losers, as well as the identities of the combatants.

#### 2.3.2 Determination of group dominance rank

Following confirmation of the linearity of the dominance hierarchy [35], the I & SI approach, which aims to determine the optimal ordering of dominance rank [36], was used to identify the dominance rankings of individuals in the experimental group. This method iteratively adjusts rank sequences to meet two criteria: (1) minimising the number of inconsistent dyads (Inconsistency, ‘I’), where the observed dominance relationship conflicts with that expected from the overall hierarchy; and (2) minimising rank distance between individuals within inconsistent dyads (strength of inconsistency, ‘SI’) [35,37]. In each dyad, the individual winning ≥ 66% of their agonistic interactions was treated as dominant, whereas the other was considered the subordinate. In dyads with just four or three agonistic interactions in daily observation data, the individual who won ≥ 75% or 100% of the interactions was considered dominant. For dyads that did not match the winning (losing) rate criteria, we considered the birds’ relationship to be tied with no dominance. If two or fewer agonistic interactions were seen in a dyad, the relationship between the two birds was deemed unknown.

A dominance relationship matrix was generated by allocating 1 and 0 to the dominant and subordinate individuals in a dyad, respectively, and 0.5 to 2 individuals with a tied relationship. We quantified the linearity of the dominance hierarchy using *h’*, an index of rank linearity [35]. The chi-square test was used to determine the statistical significance of linearity at a 1% level. Following confirmation of linearity, the dominance ranks of individuals were computed using the I & SI procedures [36]. A total of 458 agonistic interactions were recorded, with all 45 dyads showing unambiguous dominance patterns. We found significant linearity in dominance ranks across the 10 individuals (*h’* = 0.95, *χ²* = 61.26, *df* = 81, *p* < 0.01). Using the I & SI method, we determined the ranks of 10 birds with only one inconsistency between the 4^th^- and 7^th^-ranked males, whose dominance relationship was reversed concerning their ranks (the 7^th^-ranked male was dominant over the 4^th^-ranked male).

### 2.4 Experimental procedure

#### 2.4.1 Feeder box and experimental design

To examine the effect of fluctuated environments, specifically repeated changes in the spatial food distribution, on foraging performance, we conducted a group foraging experiment wherein the relationship between food amounts (none, middle, and rich) and locations was changed regularly over 3 months from October to December 2023. The experiment took place in one half of a home aviary, which was separated by a mesh door (5 × 5 × 3 m; Figure 1a and b).

**Figure 1.**
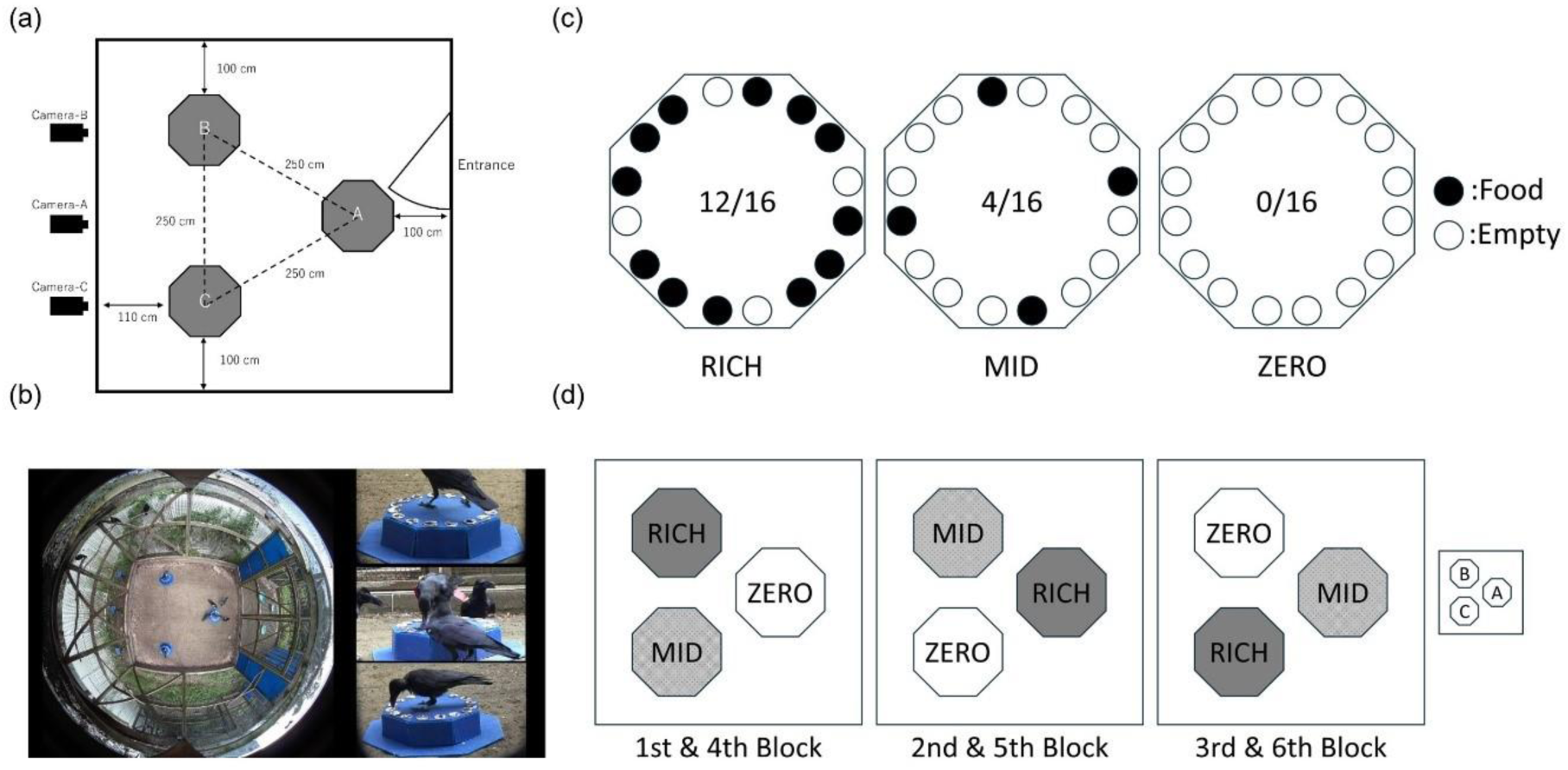
Experimental settings. (a) Feeding boxes (A, B, C) were place at three locations on the floor of the aviary. (b) Bird behaviours video recorded from above with a spherical camera and from the sides with three handy cams outside the aviary. (c) The number of cups containing food varied depending on the box condition: 12 cups in RICH, 4 cups in MID, and 0 cups in ZERO. The positions of food-containing cups in each box were pseudo-randomly varied between trials. (d) Box conditions at the three locations were maintained within blocks and moved to the next location with a clockwise direction between the blocks.

Artificial foraging patches were created using three identical plastic feeder boxes in the shape of a regular octagonal prism (side length, 16.6 cm; height, 10.0 cm) mounted on an octagonal plate (side length, 24.9 cm). At the top of each box, 16 holes (4.5 cm in diameter) were placed at 6-cm intervals to accommodate a plastic food cup. At the start of the trial, the top of each cup was covered with opaque aluminium foil to prevent the crows from seeing the interior which contained either a piece of cheese (0.3 g) or nothing (empty), depending on the box condition. The subject crows had to break the foil with their beaks to obtain food.

Each experimental trial included three feeder boxes placed 2.5 m apart on the floor of the aviary (box locations A, B, and C in Figure 1a and 1b). The boxes at these locations were identical in shape and colour, but the amount of food contained in the cups (the different conditions) varied, such as ‘RICH’, where 12 out of 16 cups contained food (Figure 1c), ‘MID’, where 4 out of the 16 cups contained food, and ‘ZERO’, where no cup contained food. In the RICH and MID conditions, the cups containing food in each box were arranged in a pseudo-random manner between trials.

The experiment included 240 trials divided into 6 blocks of 10 sessions (1 session consisted of 4 trials). We implemented two sessions (2 × 4 trials each day) from 9:00−12:30 and 13:30–16:30 and the intervals between successive daily two sessions ranged from 0 to 3 days. The relationship between food condition and box locations rotated clockwise over the blocks (Figure 1d). Crows in each block had 10 sessions (5 days × 8 trials) under stable conditions for 5 days. However, after the end of each block, the food conditions were changed for the first trial of the next session without any signals. Each trial began when the researcher (Y. N.) removed the covers from the three feeder boxes, allowing the crows to forage, and exited the experimental room. The trial was terminated 3 min after it began when the researcher removed the boxes. The inter-trial interval was approximately 20 min.

#### 2.4.2 Coded behaviour during the experiment

The behaviour of the crows during the trials was video-recorded using an omnidirectional camera (SP360-4K, Kodak) on the ceiling of the aviary and three video cameras (HD CX535, Sony, New York, NY, USA) placed outside the aviary (Figure 1b). The coded behaviours were ‘*staying on a box*’, ‘*opening a cup’*, and ‘*gaining a food*’ (Table 1). ‘Opening a cup’ was coded up to a maximum of 48 times per trial (3 boxes × 16 cups). Crows that investigated an already-opened cup were not considered. ‘Gaining a food’ was recorded up to 16 times (RICH: 12 cups, MID: 4 cups). We recorded the individual identity and timing of these actions. The time resolution for behavioural coding was approximately 0.3 s based on the video frame. The software BORIS v. 7 [34] was used for behaviour coding.

**Table 1.**
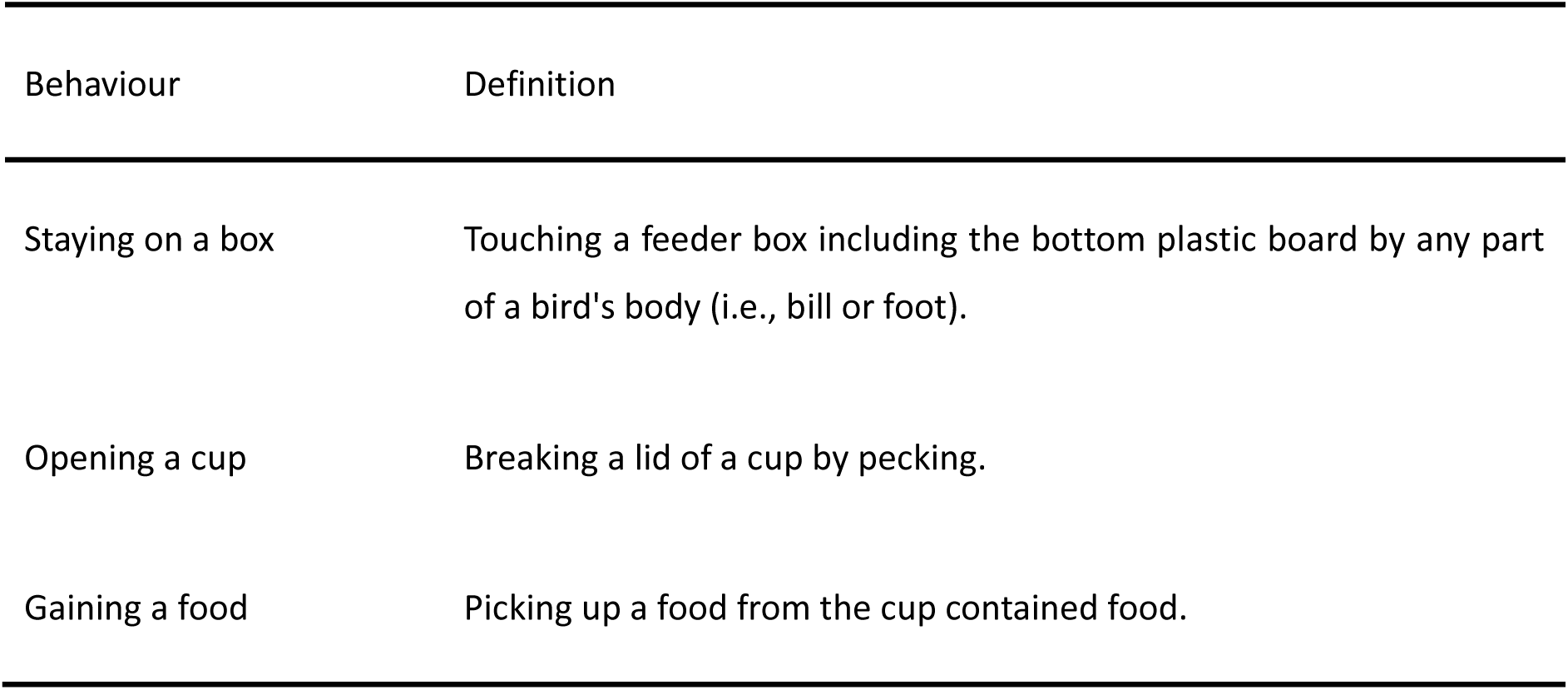
Coded behaviours.

| Behaviour | Definition |
| --- | --- |
| Staying on a box | Touching a feeder box including the bottom plastic board by any part of a bird's body (i.e., bill or foot). |
| Opening a cup | Breaking a lid of a cup by pecking. |
| Gaining a food | Picking up a food from the cup contained food. |

### 2.5 Analysis

Based on the observed behaviours, we derived indices of group foraging for each trial, such as ‘*number of successful foragers’,* ‘*foraging duration’,* and ‘*exploring duration’,* to evaluate foraging efficiency at the group level (Table 2). The ‘number of successful foragers’ was defined as the number of crows that obtained at least one food item (one piece of cheese) in a trial. ‘Foraging duration’ was defined as the time between the first ‘opening a box’ and the final (16^th^) event of ‘gaining a food’, regardless of box condition or individual. ‘Exploring duration’ was defined as the time between the first ‘staying on a box’ event and the 48^th^ ‘opening a cup’. In circumstances where not all 48 cups were opened for 3 min, we calculated the time from the initial ‘staying a box’ event to the last ‘opening a cup’ event. In addition, we estimated the Gini coefficient to evaluate the degree of inequality in food gain distribution among individuals. The Gini coefficient was determined using a Lorenz curve [38] derived from the cumulative ratio of subjects to food gains, which ranged from 0 (not skewed) to 1 (highly skewed). We calculated and adjusted the Gini coefficient to account for the small sample size using the ‘ineq’ R package [39]. Out of 240 trials, 18 were excluded from the analysis: 1 due to video recording issues, 1 where not all the food was consumed, and 16 from the 7^th^ to 10^th^ sessions of the 6^th^ block due to the removal of an individual for health issues.

**Table 2.** Indices of group foraging.

| Index | Definition |
| --- | --- |
| Gini coefficient | A measure of inequality in food-gain distribution, calculated from the Lorenz curve [39] obtained from the cumulative ratio of the number of subjects and food gains. This index ranged between 0 and 1. |
| Number of successful foragers | The number of individuals that gained one or more food(s) in a trial. |
| Foraging duration | The duration from the first 'opening a cup' to the last 'gaining a food' event in a trial. |
| Exploring duration | The duration from the onset of the first 'staying on a box' event to the last 'opening a cup' in a trial. |

#### 2.5.1 Group-level analysis

We used generalised linear models (GLMs) to examine the effects of changing environment on group foraging. We performed GLMs independently on group foraging indices as response variables, namely the Gini coefficient (with Gaussian error distribution and identity link function), the number of successful foragers (with Poisson error distribution and log link function), foraging duration (with Gamma error distribution and log link function), and exploration duration (with Gamma error distribution and log link function). Furthermore, for each response variable, we performed two lines of model analysis at distinct timescales: short- and long-term effects. The short-term scale analysis examined the immediate effects of environmental change on each foraging index between sessions; thus, GLMs included a session component as a fixed effect. For this short-term analysis, we divided the sessions into three types: ‘*pre-change session’,* which included the 10^th^ (final) sessions of blocks as the data just before the environmental change; ‘*post-change sessio*n’, which included the first sessions of blocks as the data just after the environmental change; ‘*post-change + 1 session’,* which included the second sessions of blocks as the data following the post-change session. The long-term scale analysis, on the contrary, investigated the cumulative effects of repeated environmental changes on each foraging index between blocks; therefore, GLMs included a block factor as a fixed effect.

#### 2.5.2 Individual-level analysis

The effects of environmental changes and individual social characteristics, such as dominance rank and sex, on foraging performance were examined. We used GLMs, with the number of food gains per individual as the response variable, session number, block number, dominance rank, sex, and their secondary interactions as independent variables with negative binomial error distribution and log link function. To address multicollinearity, rank and sex were not included in the same model. Instead, in the individual-level model selection, the best model was chosen from those that included either rank or sex.

All model analyses were conducted using the statistical software R v. 4.3.2 [40]. Before constructing the models, we checked for overdispersion of the response variable using the ‘performance’ package [41]. We used Akaike’s Information Criterion (AIC), based on smaller is better, to select the best model. Model selection was performed using the ‘MuMIn’ package [42] and probabilities and confidence intervals were estimated using the ‘ggeffect’ program [43] with a significance level of 0.05.

## 3. Results

### 3.1 Group level

#### 3.1.1 Short-term effects

GLM analysis of the Gini coefficients (a measure of inequality ranging from 0 to 1, with values closer to 1 indicating greater inequality; Table 2) revealed a significant effect of the post-change session compared with the pre-session, with a positive coefficient indicating an immediate increase in the Gini coefficient following the environmental change (AIC_best(session)_ = −117.266, AIC_null_ = −97.262; Table 3 and Figure 2a). This result demonstrates that environmental changes immediately increase inequality in food gain among individuals.

**Figure 2.**
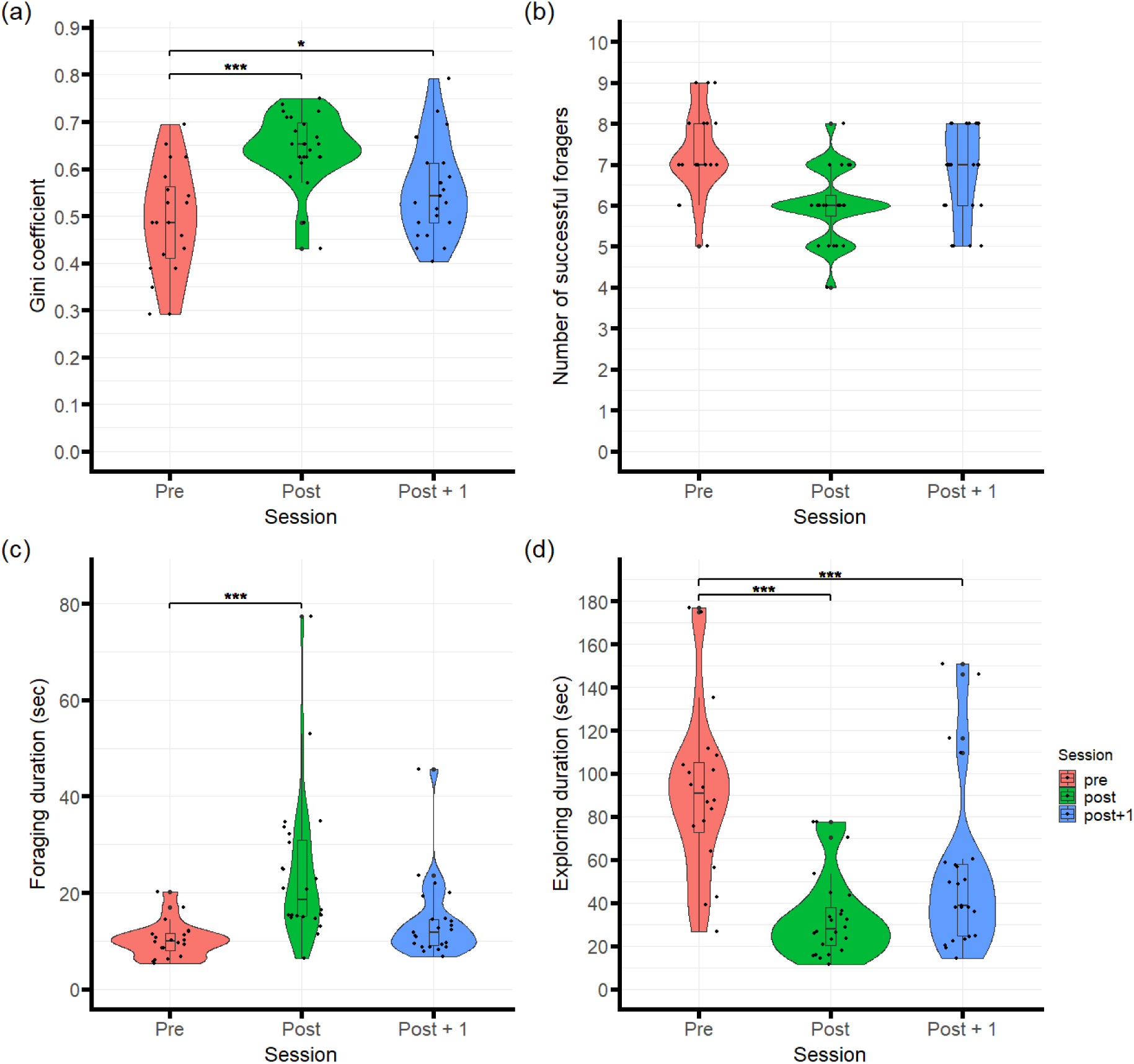
Short-term changes after environmental fluctuations. (a) The inequality of food distribution increased. (b) No short-term effect was observed in the number of successful foragers. (c) The time required to consume all foods at each trial increased. (d) The time to open cups, including the ZERO condition box (without food), decreased. Pre: the 10^th^ session of each block (immediately before fluctuations). Post: first session of each block (just after changes). Post + 1: second session of each block. Plots show the data at each trial. Asterisks indicate significant differences (\**p* < 0.05, \*\**p* < 0.01, \*\*\**p* < 0.001).

**Table 3.** Outputs of the best model from a GLMs in short-term effect analysis.

| Index | Variables | Estimate | S.E. | Test Statistic | *p* |
| --- | --- | --- | --- | --- | --- |
| Gini coefficient | **Intercept** | **0.490** | **0.022** | ***t* = 22.555** | **< .001** |
|  | **Session** |  |  |  |  |
|  | **Post** | **0.153** | **0.029** | ***t* = 5.207** | **< .001** |
|  | **Post + 1** | **0.066** | **0.030** | ***t* = 2.237** | **0.029** |
| Number of successful foragers | **Intercept** | **1.898** | **0.047** | ***z* = 40.125** | **< .001** |
| Foraging duration | **Intercept** | **2.328** | **0.124** | ***t* = 18.819** | **< .001** |
|  | **Session** |  |  |  |  |
|  | **Post** | **0.855** | **0.167** | ***t* = 5.103** | **< .001** |
|  | Post + 1 | 0.314 | 0.169 | *t* = 1.854 | 0.068 |
| Exploring duration | **Intercept** | **4.523** | **0.133** | ***t* = 34.004** | **< .001** |
|  | **Session** |  |  |  |  |
|  | **Post** | **-1.014** | **0.180** | ***t* = -5.629** | **< .001** |
|  | **Post + 1** | **-0.535** | **0.182** | ***t* = -2.940** | **0.005** |

The 'Session' term compared the sessions just before the environmental change ('Post' session: the 10<sup>th</sup> sessions of blocks) with the session after the changes (Post: the 1<sup>st</sup> sessions of blocks) and the session after that (Post + 1: the 2<sup>nd</sup> sessions of blocks). Gini coefficient was analysed Gaussian error distribution and identity link function. The number of successful foragers was analysed using Poisson error distribution and log link function. Foraging duration and exploration duration were analysed using Gamma error distribution and log link function. Bold indicates statistically significant results.

For the number of successful foragers (the number of crows that obtained at least one food item; Table 2), GLM analysis indicated that the null model was the best, showing that environmental change had no short-term effect on the number of successful foragers (AIC_best(null)_ = 265.905, AIC_session model_ = 266.995; Table 3 and Figure 2b).

The best GLM model for foraging duration (the time taken to consume all the food; Table 2) showed a significant effect of the post-change session compared to the pre-change session, with a positive coefficient, indicating that foraging duration increased immediately after the environmental change (AIC_best(session)_ = 451.164, AIC_null_ = 478.993; Table 3 and Figure 2c).

For exploration duration (the time taken to explore for food around the feed boxes; Table 2), the best model produced a significant negative coefficient for the post-change session compared with the pre-change session, indicating decreased in exploration duration immediately after the environmental change (AIC_best(session)_ = 634.046, AIC_null_ = 661.451; Table 3 and Figure 2d).

#### 3.1.2 Long-term effects

GLM analysis of the Gini coefficient identified the best model as having a significant negative correlation with the *session × block* interaction, but no significant correlation with *session* or *block* individually (AIC_best_ = −450.861, AIC_null_ = −386.852; Table 4 and Figure 3a). This result indicates that the Gini coefficient declined progressively throughout the sessions, with effects intensifying as blocks progressed, demonstrating that repeated environmental changes reduced inequality in food gains among individuals over time.

**Figure 3.**
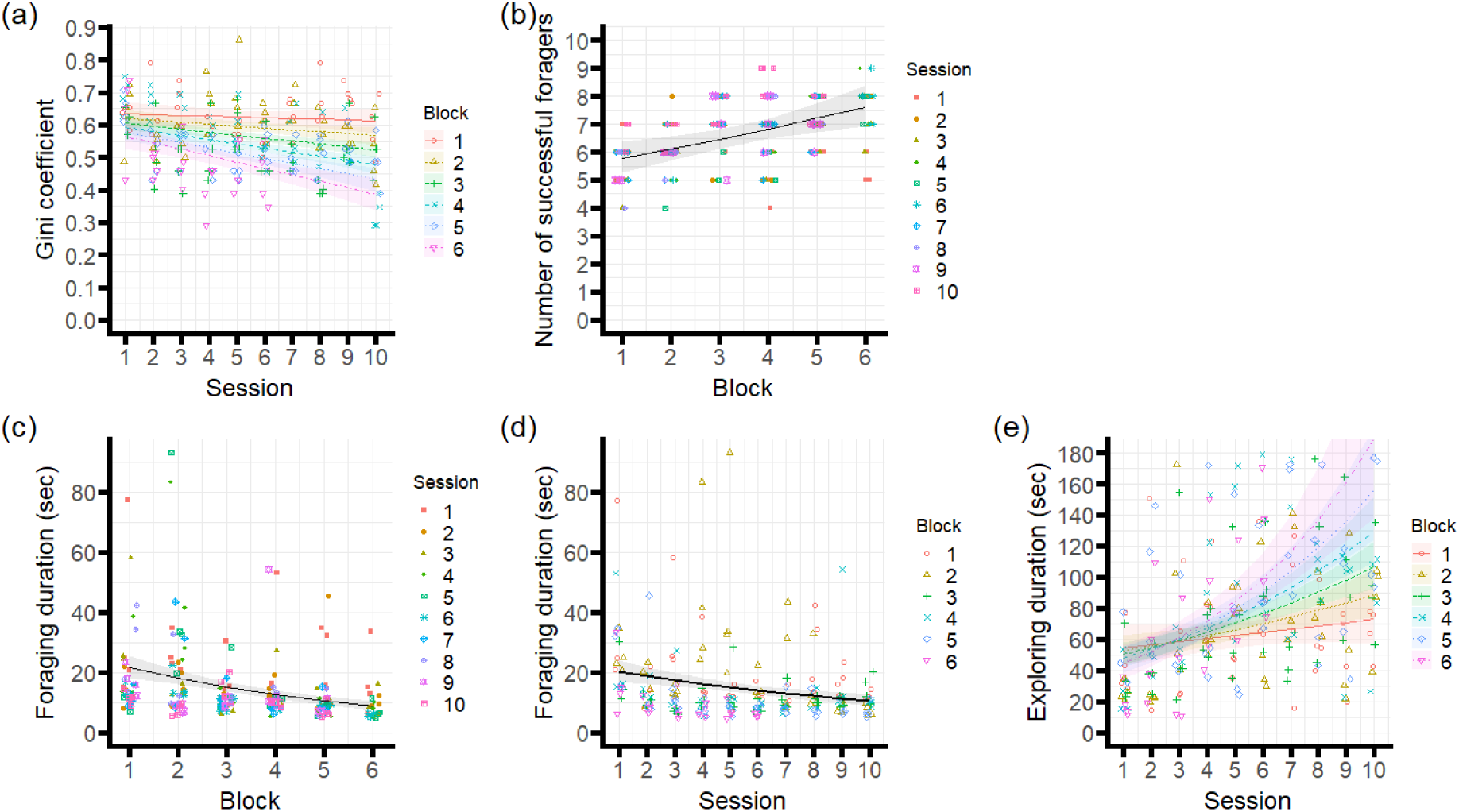
Long-term changes after environmental fluctuations. (a) The reduction of inequality in food distribution with session advancement became more pronounced as the blocks progressed. (b) As the blocks progressed, so did the number of successful foragers, with more individuals acquiring at least one food at each attempt. (c) The duration of foraging was reduced as the experiment advanced. (d) The time to consume all foods at each trial decreased as the sessions progressed. (e) The time to open cups increased as the session progressed. This effect became more marked with the passage of blocks. Plots show the data at each trial with colours indicating (a)(d)(e) blocks and (b)(c) sessions. The regression lines show the predicted value of the best model. The shaded areas around the lines indicate a 95% confidence interval.

**Table 4.** Outputs of the best model from a GLMs in long-term effect analysis.

| Index | Variables | Estimate | S.E. | Test Statistic | *p* |
| --- | --- | --- | --- | --- | --- |
| Gini coefficient |  |  |  |  |  |
|  | **Intercept** | **0.647** | **0.028** | ***t* = 22.839** | **< .001** |
|  | Session | 0.001 | 0.005 | *t* = 0.223 | 0.824 |
|  | Block | -0.010 | 0.007 | *t* = -1.401 | 0.163 |
|  | **Session × Block** | **-0.003** | **0.001** | ***t* = -2.635** | **0.009** |
| Number of successful foragers |  |  |  |  |  |
|  | **Intercept** | **1.632** | **0.085** | ***z* = 19.304** | **< .001** |
|  | Session | 0.014 | 0.009 | *z* = 1.502 | 0.133 |
|  | **Block** | **0.055** | **0.016** | ***z* = 3.370** | **< .001** |
| Foraging duration |  |  |  |  |  |
|  | **Intercept** | **3.610** | **0.148** | ***t* = 24.433** | **< .001** |
|  | **Session** | **-0.071** | **0.029** | ***t* = -4.244** | **< .001** |
|  | **Block** | **-0.178** | **0.029** | ***t* = -6.110** | **< .001** |
| Exploring duration |  |  |  |  |  |
|  | **Intercept** | **4.041** | **0.180** | ***t* = 22.481** | **< .001** |
|  | Session | 0.006 | 0.030 | *t* = 0.206 | 0.837 |
|  | Block | -0.068 | 0.047 | *t* = -1.452 | 0.148 |
|  | **Session × Block** | **0.026** | **0.008** | ***t* = 3.087** | **0.002** |

The number of successful foragers was analysed using Poisson error distribution and log link function. Foraging duration and exploration duration were analysed using Gamma error distribution and log link function. The Gini coefficient was analysed Gaussian error distribution and identity link function. Bold indicates statistically significant results.

GLM analysis of the number of successful foragers over time yielded a best model that included session and *block* variables (AIC_best model_ =865.462; AIC_null_ = 873.937). The best model showed no significant differences across *session;* however, a significant positive effect of *block* was observed (Table 4 and Figure 3b), indicating that the number of successful foragers increased across experimental blocks.

For foraging duration, the best GLM model showed significant negative effects of both *session* and *block* (AIC_best model_ = 1474.716, AIC_null_ = 1548.145; Table 4 and Figure 3c and 3d), indicating that foraging time decreased across sessions and blocks.

GLM analysis revealed a significant positive correlation between the exploration duration and session × block interaction, with no significant effects for either session or block (AIC_best model_ = 2220.574, AIC_null_ = 2265.159; Table 4 and Figure 3e). This result shows that exploration duration increased throughout sessions within blocks, with effects increased as the blocks progressed.

### 3.2 Individual level

GLM analysis examining the influence of dominance rank or sex on food gain identified the best model as having a significant negative correlation with *block* × *sex* but no significant correlation with either *sex* or *block* (AIC_best_ = 7549.073, AIC_null_ = 7560.038; Tables 5 and 6; Figures 4 and S1). Stratified GLMs applied separately on male and female datasets produced the best model for males (AIC_best(block)_ = 3712.150, AIC_null_ = 3712.712), and females (AIC_best(block)_ = 3787.882, AIC_null_ = 3789.328; Table S1), both of which included blocks but showed no significant correlations. These findings indicate that food acquisition decreased in males but increased in females over time, with females gaining more food than males in later experimental blocks.

**Figure 4.**
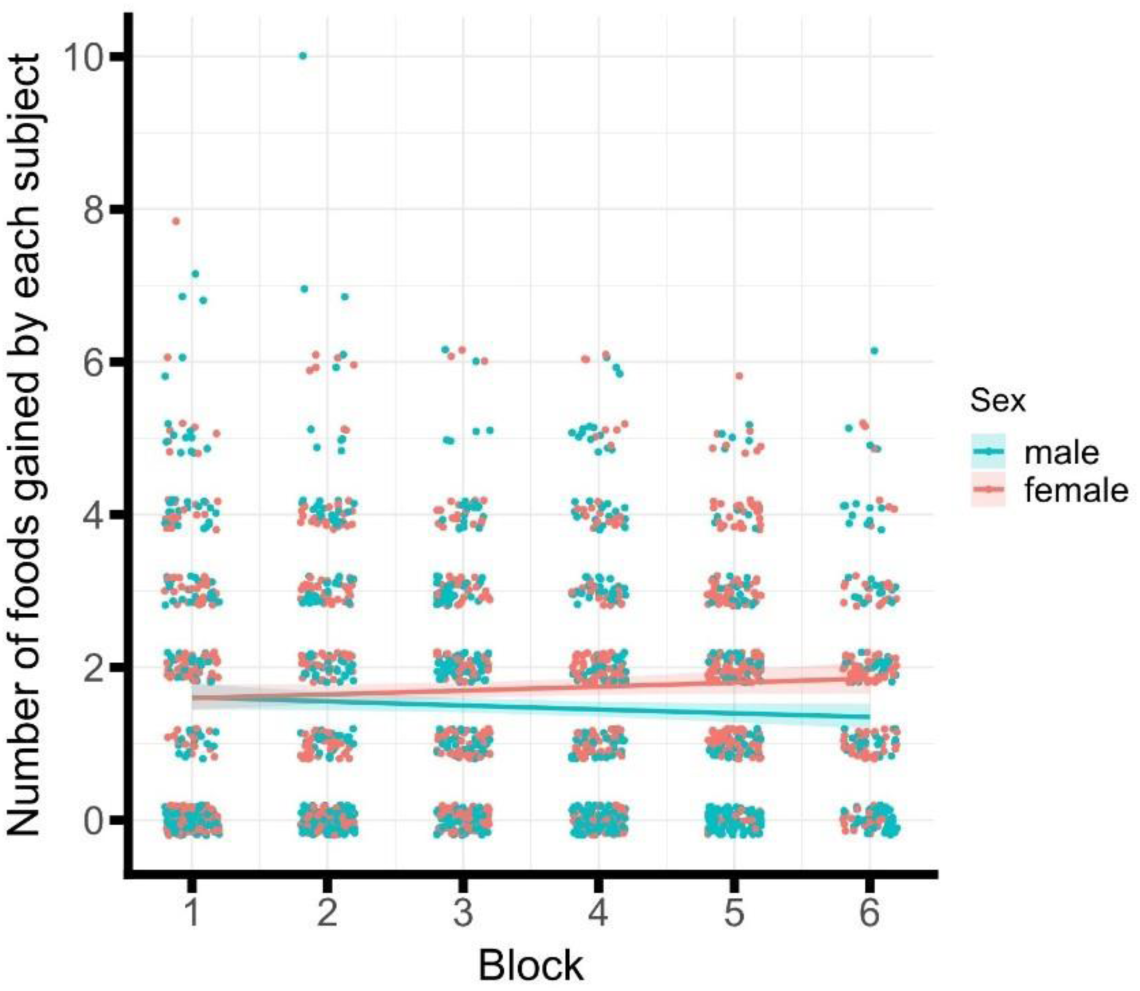
Effects of subjects’ sex on the number of food items acquired by each subject. As the blocks progressed, the amount of food consumed by females surpassed that of males. The jitter plots show the data of each subject by trial. Regression lines show the predicted values of the best model. Shaded areas around the lines indicate a 95% confidence interval.

**Table 5.** Result of model selection in individual level.

| Model | Model Formula | AIC | ΔAIC | df |
| --- | --- | --- | --- | --- |
| **Best Model of Sex Model** | **Intercept + Block + Sex + Block×Sex** | **7549.073** | - | **5** |
| Full Model of Sex Model | Intercept + Block + Session + Sex +<br>Block×Session + Block×Sex +<br>Session×Sex + Block×Session×Sex | 7554.709 | 5.636 | 9 |
| Best Model of Rank Model<br>(Null Model) | Intercept | 7560.038 | 10.96<br>5 | 2 |
| Full Model of Rank Model | Intercept + Block + Session + Rank+<br>Block×Session + Block×Rank+<br>Session×Rank+ Block×Session×Rank | 7571.504 | 22.43<br>1 | 9 |

Each model was analysed using negative binomial error distribution and log link function. Bold indicates the best model of individual level models.

**Table 6.**
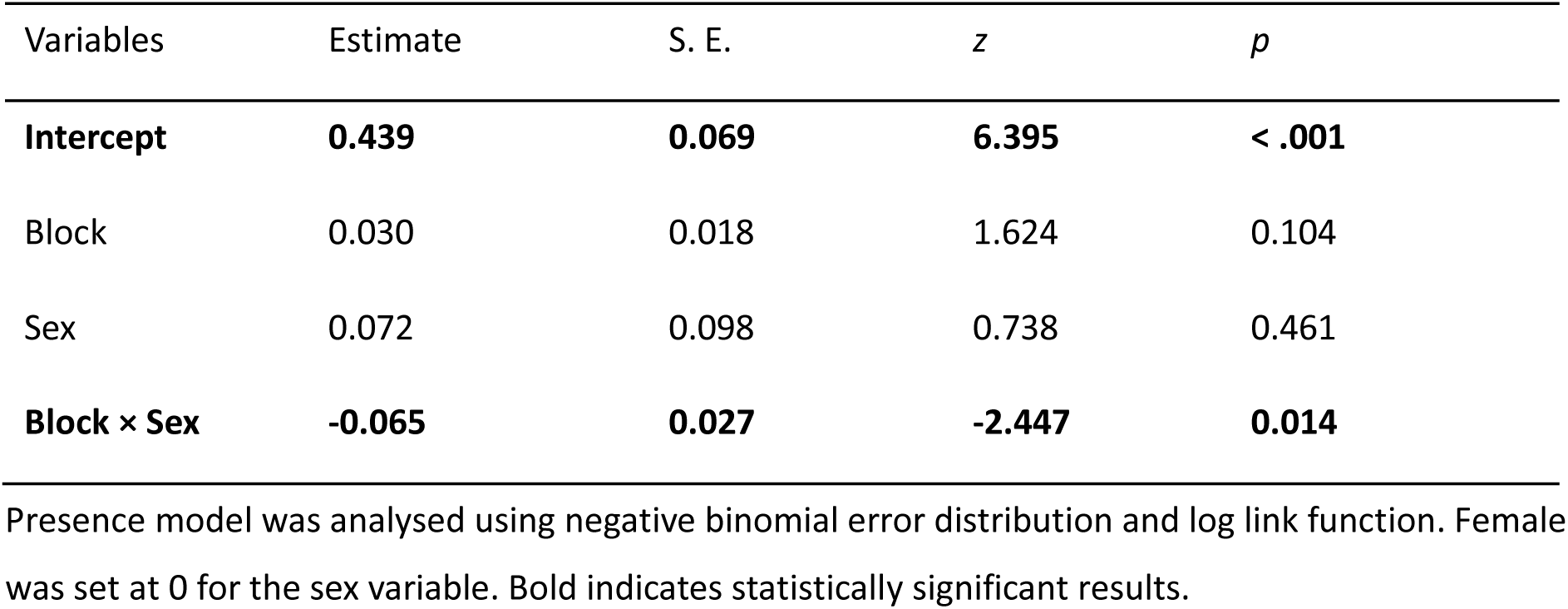
Outputs of the best model from a GLM testing the effect of the sex on food gain.

| Variables | Estimate | S. E. | *z* | *p* |
| --- | --- | --- | --- | --- |
| **Intercept** | **0.439** | **0.069** | **6.395** | **< .001** |
| Block | 0.030 | 0.018 | 1.624 | 0.104 |
| Sex | 0.072 | 0.098 | 0.738 | 0.461 |
| **Block × Sex** | **-0.065** | **0.027** | **-2.447** | **0.014** |

Presence model was analysed using negative binomial error distribution and log link function. Female was set at 0 for the sex variable. Bold indicates statistically significant results.

## 4. Discussion

In this study, we experimentally examined how a fluctuating environment—specifically, repeated changes in the relationship between food quantity and spatial location—affected inequality of food acquisition and foraging performance within captive group of large-billed crows. Additionally, we investigated the influence of dominance rank and sex on the individual food acquisition patterns during environmental change.

The observed changes in the Gini coefficient showed a gradual reduction in food acquisition inequality over long-term. This result appears to be driven by females, who gradually increased the food gain in the later blocks of the experiment. In large-billed crows, the frequency of aggressive interactions from males towards females has been reported to be lower than that among males, and aggression among females occurs even less frequently [31]. The differences in the frequency of aggression may cause low pressure to displace the females from the rich or middle food sites, contributing to this result.

In this experiment, environmental conditions were regularly altered, creating a challenging situation in which individuals were required to adjust their previously learned behaviour accordingly. Individuals with higher aggression frequencies performed more poorly on associative learning tasks, and it has been suggested that this may be due to a reduced allocation of time and energy to attend to the task [26]. This raises the possibility that males, who were generally more aggressive than females, may have been less able to cope with the changing environment.

Moreover, the total amount of food was limited to fewer than two items per individual in this experiment, thus necessitating rapid foraging. As overall foraging efficiency increased across sessions, all food was typically consumed within the first several seconds. Under such conditions, from the perspective of food defensibility [19], initiating aggression to monopolies food as well as being targeted by aggression may have had negative consequences for food intake. In contrast, females— who were less likely to be disrupted by others—may have been able to develop their own optimal foraging strategies under relatively high levels of uncertainty [27]. In addition, sex differences in the use of social information [6] and in motivation for foods [33] may also have conferred advantages under uncertain conditions [28]. These findings suggest that environmental change can promote alternative foraging strategies and opportunities not observed under stable conditions, ultimately contributing to the reduction of inequality in food acquisition.

The increase in the Gini coefficient immediately after each environmental change indicates that some individuals successfully adapted to the new conditions and temporarily acquired a larger share of food. However, contrary to our initial expectation, we did not observe a consistent pattern where subordinate individuals flexibly switched foraging sites and obtained more food in the short-term. Rather, the changes in food acquisition following environmental fluctuations varied across individuals (Fig S1). These findings suggest that individuals’ social characteristics such as dominance rank may only partially explain their short-term foraging behaviour. Some studies have highlighted the importance of social networks in social foraging (e.g., [20,44]), indicating the need for more detailed examinations of social interactions. Understanding who forages with whom and whose information is used may be crucial, especially considering that such social networks have been regarded as one important component of wealth (‘relational wealth’) [1,3].

Our experimental manipulation altered the external foraging environment while maintaining constant group composition. This approach contrasts with those used in previous research on zebra finches (*Taeniopygia guttata*), which involved repeatedly dividing stable groups and introducing short-term disruption through group reunification [45]. The study’s manipulation of the internal social environment resulted in recurrent short-term changes and long-term decline in foraging efficiency, contrasting with our findings. In our study, the stable group composition may have facilitated consistent use of social information from conspecifics. Individual unable to adapt to environmental changes independently may have relied on information from more adaptable group members, thereby maintaining overall group foraging efficiency. These results suggest the importance of maintaining stable social structures even at the potential cost of individual equality, particularly in environments characterised by frequent change. The ability to maintain group cohesion whilst adapting to environmental variability may represent a key advantage of social foraging systems.

Repeated short-term environmental changes caused a gradual decrease in the Gini coefficient. As frequent and intense climatic changes were likely to prevent resource monopolisation in early human societies [2], short-term environmental changes may reduce social inequality. The similarity between this and our results suggests that common mechanisms may contribute to the formation of inequality across humans and other animal societies. Future research on group foraging should consider both internal and external social factors. This comprehensive approach may lead to a deeper understanding of social animal groupings and its dynamics.

## 5. Conclusion

Overall, our finding demonstrates a gradual reduction in food acquisition inequality over the long-term, as evidenced by declining the Gini coefficients across experimental blocks. This pattern appears driven primarily by females, who progressively increased their food gain in later blocks of the experiment. This sex-specific response may be explained by differences in aggressive behaviour patterns within large-billed crows. These results highlight the importance of incorporating environmental changings into studies of group foraging. Rather than viewing social dominance as a fixed determinant of foraging success, our findings suggest that environmental context fundamentally shapes the expression and consequences of social relationships. Future group foraging research should adopt comprehensive approaches that consider both internal social factors and external environmental fluctuations to better understand the complex dynamics governing resource acquisition in animal societies.

The methodological approach employed in this study provides a valuable framework for future investigations into the mechanisms underlying inequality in animal societies, including humans. Such research may ultimately contribute to deeper understanding of the processes that generate and maintain social inequality across diverse biological and social systems.

## Acknowledgments

We are grateful to Aymi Motai and Kosuke Miyazaki for conducting prototype experiments of the present study. We also thank the members of the Hiyoshi field station of Animal Psychology Lab, Keio University for their support in the daily care of the animals.

## Funding Statement

This study was supported for research of this work from JSPS [KAKENHI grant numbers 21H04421, 21H00962, 23K25755, 23H00074] and Keio University Grant-in-Aid for Innovative Collaborative Research Projects [Grant number MKJ1905].

## Data Accessibility

The data generated and analysed during this study are available from https://doi.org/10.6084/m9.figshare.27124326.v3

Example videos of behavioural experiments are also available from https://doi.org/10.6084/m9.figshare.27124350.v1 (pre-session), https://doi.org/10.6084/m9.figshare.27124344.v2 (post-session), https://doi.org/10.6084/m9.figshare.27124356.v1 (post + 1 - session).

The R script with all the code used during this study is available in https://github.com/2ki2ki/crow_RepEnvChange/tree/main.

## Competing Interests

We declare we have no competing interests.

## Authors’ Contributions

Y.N.: Writing – original draft, Visualisation, Validation, Methodology, Investigation, Formal analysis, Data curation, Conceptualisation.

M.T.: Validation, Formal analysis, Investigation, Data curation.

E-I.I.: Writing – review & editing, Resources, Validation, Supervision, Project administration, Methodology, Investigation, Data curation, Funding acquisition, Conceptualisation.

## Notes

https://doi.org/10.6084/m9.figshare.27124326.v3

https://doi.org/10.6084/m9.figshare.27124350.v1

https://doi.org/10.6084/m9.figshare.27124344.v2

https://doi.org/10.6084/m9.figshare.27124356.v1

https://github.com/2ki2ki/crow_RepEnvChange/tree/main

### Competing Interest Statement

The authors have declared no competing interest.

### Summary of Updates

This version has been revised to refine the Introduction and the Discussion, improving the clarity and precision of the main arguments.

