## Supplemental Table 1 and Figure 1 for "Repeated environmental changes mitigate inter-individual inequality of food gain in a group foraging experiment with large-billed crows"

### Supplementary Materials

Table S1. Stratified GLMs separate for the male and the female datasets

| Variables | Estimate | S. E. | *z* | *p* |
|---|---|---|---|---|
| *Males* | | | | |
| Intercept | **0.512** | **0.080** | **6.383** | **< .001** |
| Block | -0.035 | 0.022 | -1.614 | 0.107 |
| *Females* | | | | |
| Intercept | **0.438** | **0.061** | **7.239** | **< .001** |
| Block | 0.030 | 0.016 | 1.853 | 0.064 |

Presence model was analysed using negative binomial error distribution and log link function. Bold indicates statistically significant results.

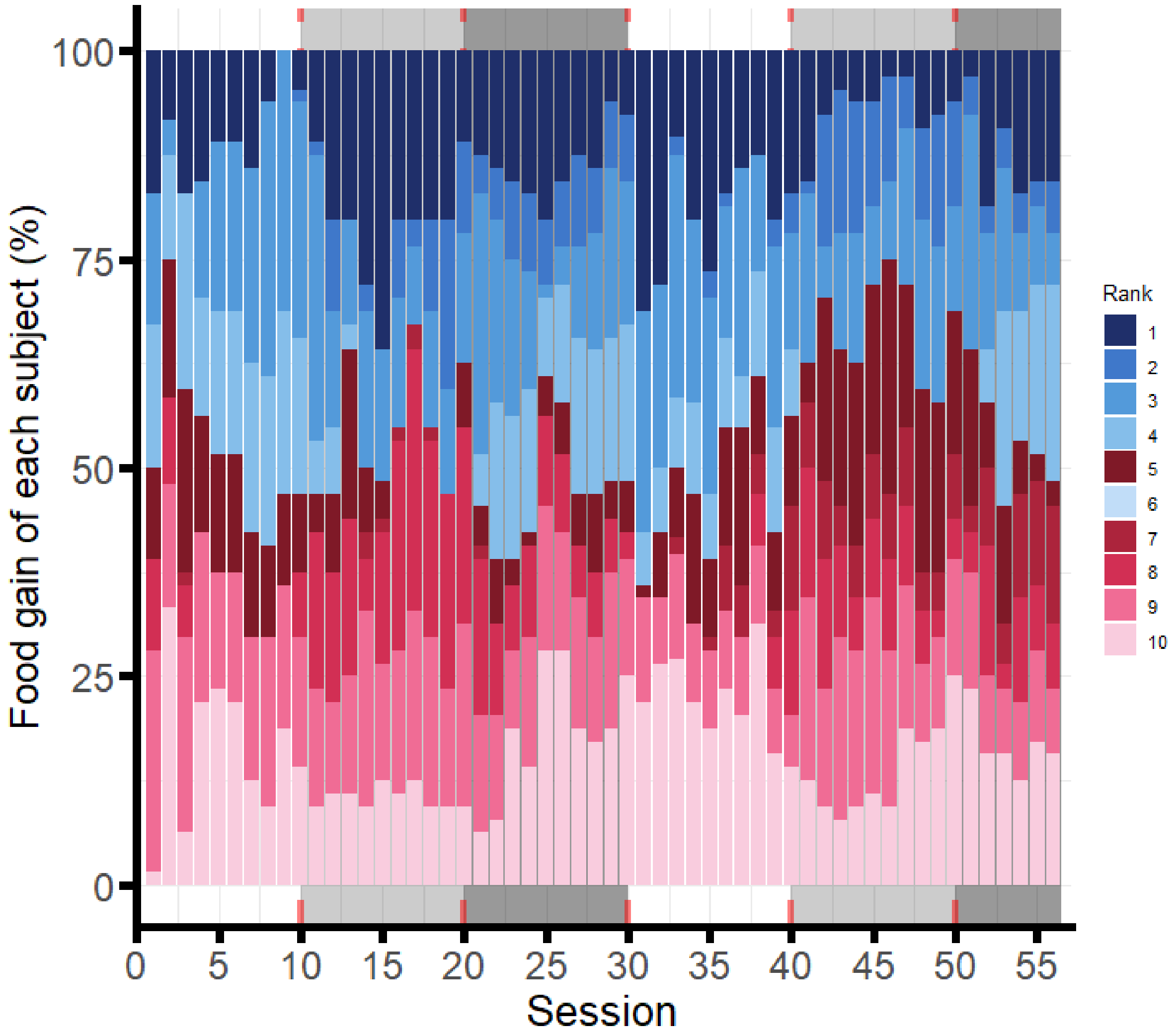


**Figure S1**. Proportion of food gain per individual across sessions. Male and female individuals are shown in blue and red, with darker shades representing higher dominance ranks. Background colours (white, grey, and black) mark sessions with identical relationships between food amount and box location.
